# The influence of heteroresistance, growth and antibiotic selection in shaping the invasion dynamics of colistin resistant *Pseudomonas aeruginosa*

**DOI:** 10.64898/2025.12.15.694340

**Authors:** Angie C. Alarcon Rios, Anh Duc Pham, Catherijne A.J. Knibbe, Daniel E. Rozen, J. G. Coen van Hasselt, Linda B.S. Aulin

## Abstract

Heteroresistance, where a small subpopulation of phenotypically resistant cells coexists within an otherwise susceptible population, plays a critical role in bacterial survival during antibiotic exposure. Yet, its influence on the invasion success of genetically resistant strains remains poorly understood. In this study, we investigate the influence of bacterial heteroresistance, growth, and antibiotic selection on the outcome of an invasion experiment. We quantified the invasion dynamics of a bioluminescent colistin-resistant *Pseudomonas aeruginosa* strain across a colistin concentration gradient during co-culture with colistin-susceptible clinical *P. aeruginosa* isolates with varying levels of heteroresistance. The observed variation in heteroresistance and fitness of the clinical isolates allowed investigation of the impact of these factors on invasion dynamics. We hypothesized that fitter isolates would limit invasion through competitive exclusion, while heteroresistant isolates, despite their growth costs, may serve as reservoirs for resistance evolution under antibiotic selection. Our results show that in antibiotic-free conditions, faster-growing isolates competitively excluded the colistin resistant invader while isolates with high levels of heteroresistance failed to do so. This competitive landscape shifted with increasing colistin concentration, giving the invader a fitness advantage that peaked around 2-4xMIC of the clinical isolate. The shift in the landscape over the colistin gradient might further have been influenced by the presence of heteroresistance in the clinical isolates. These findings reveal how both heteroresistance and competitive exclusion shapes invasion dynamics. Understanding these interactions is critical for redesigning treatment strategies that minimize ecological opportunities for resistant strains to establish and expand.

## Introduction

Heteroresistance (HR) is a bacterial survival strategy in which very small subpopulations exhibit significantly reduced antibiotic susceptibility compared to the overall population detected by standard diagnostic tests(Andersson, Nicoloff, and Hjort 2019; Pereira et al. 2021). Although often transient and unstable, HR can allow these less susceptible subpopulations to survive antibiotic selection and subsequently regrow. This strategy potentially contributes to treatment failure, antibiotic resistance development, and persistent infections(Frimodt-Møller et al. 2018; Rossi et al. 2020; Victor I. Band et al. 2018; He et al. 2018; Dewachter, Fauvart, and Michiels 2019), highlighting its clinical importance.

Despite increasing recognition of its clinical relevance, HR remains poorly integrated into diagnostic and treatment frameworks. Conventional susceptibility testing methods frequently fail to detect HR(Band et al. 2021; V I Band et al. 2018; Howard-Anderson et al. 2022), potentially underestimating the likelihood of therapeutic failure in infections classified as susceptible by MIC alone(Pereira et al. 2021). These diagnostic limitations are especially concerning amid rising antimicrobial resistance, where phenotypic heterogeneity may conceal early resistance emergence and interfere with selection dynamics.

Heteroresistance has been documented across multiple bacterial species and antibiotic classes, including last-resort agents such as colistin(Hermes et al. 2013; Band et al. 2021; 2019). Colistin is often used to treat multidrug-resistant infections caused by *Pseudomonas aeruginosa*(Falagas, Kasiakou, and Saravolatz 2005), a highly adaptable opportunistic pathogen, frequently associated with antibiotic-resistant infections in the lungs of cystic fibrosis patients(Ciofu et al. 2012; Rossi et al. 2021; Rossi et al. 2020) and severe infections in immunocompromised hospital patients worldwide(Markou and Apidianakis 2014; Pachori, Gothalwal, and Gandhi 2019; Potron, Poirel, and Nordmann 2015; Peng et al. 2014). However, the clinical efficacy of colistin is increasingly compromised by both stable genetic resistance and phenotypic survival strategies such as HR(Andersson, Nicoloff, and Hjort 2019; Gogry et al. 2021; Dößelmann et al. 2017). In *P. aeruginosa*, HR has been linked to subpopulations capable of surviving otherwise lethal antibiotic concentrations(Jie et al. 2019; Howard-Anderson et al. 2022; Hawley, Murray, and Jorgensen 2008). Although different molecular mechanisms underlying colistin HR have been identified(Andersson, Nicoloff, and Hjort 2019; Hjort, Nicoloff, and Andersson 2016), its ecological and evolutionary implications remain poorly understood.

One ecological and evolutionary aspect that is currently under-investigated is the influence of HR on invasion dynamics, which relates to the introduction of a resistant invader strain into a larger sensitive bacterial population. While antibiotic selection pressure is traditionally viewed as a directional force that eliminates susceptible bacteria and favors resistant variants(Holmes et al. 2016; Hughes and Andersson 2017; MacLean and San Millan 2019), the fixation dynamics of resistant strains can be more complex than simple population replacement of susceptible populations by resistant clones. In otherwise susceptible strains where the overall population is kill by the antibitic, heteroresistant subpopulations may act as bacterial reservoirs that can acquire *de novo* resistance and take over as a new resistant population (Hughes and Andersson 2017; MacLean and San Millan 2019; Xu et al. 2025; Mallon, Van Elsas, and Salles 2015). Conversely, these heteroresistant sub-populations may delay the population replacement by fully resistant clones by imposing competitive and resource constraints that limit their establishment. Such effects are likely to be most pronounced at sub-lethal antibiotic concentrations(Andersson and Hughes 2012), where the fitness trade-offs, i.e. lower growth rates, often associated with resistance and HR can shape population dynamics and invasion potential(Levin, Perrot, and Walker 2000; Lenski et al. 2015).

To address the knowledge gaps related to HR and invasion dynamics, this study investigates how HR and growth influences the invasion dynamics of a colistin-resistant *P. aeruginosa* strain.

## Materials and Methods

This study implemented an experimental approach to investigate the invasion dynamics of a focal colistin-resistant strain of *P. aeruginosa* into different colistin-susceptible clinical isolates with varying levels of HR. The workflow is summarized in **Figure 1**.

**Figure 1.**
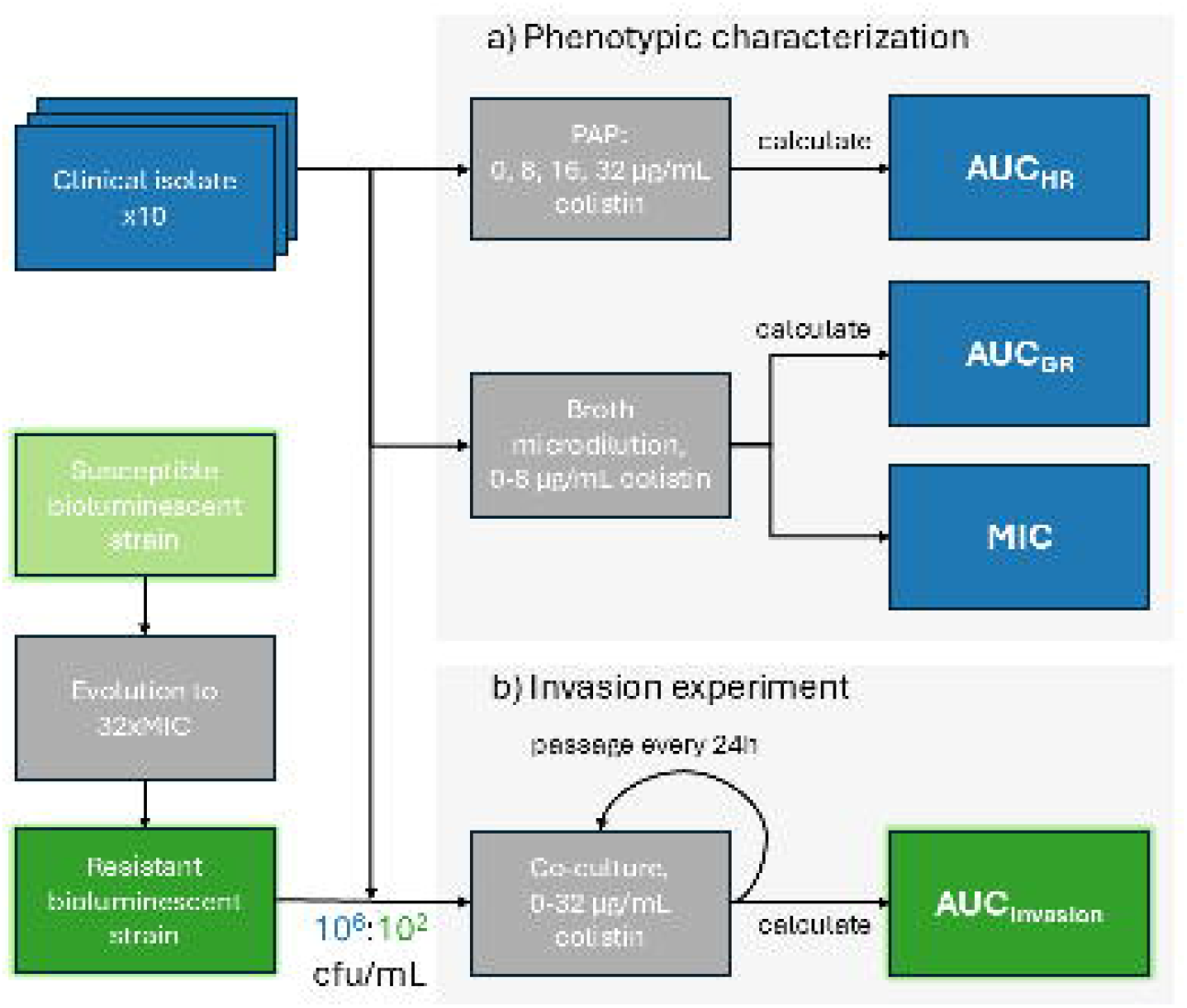
Experimental workflow. (a) Phenotypic characterization of clinical isolates to determine growth (OD_600_), minimum inhibitory concentration (MIC) using broth microdilution, as well as population analysis profile (PAP) tests to determine heteroresistance (HR). (b) Invasion-experiment where the resistant invader strain is co-cultured with each clinical isolate at an initial ratio of 10^2^ CFU/mL (invader) to 10 □ CFU/mL (clinical isolate). Cultures were maintained in 200µl 96-well plates with serial passaging every 24 hours. Luminescence relative light units from the invader strain were recorded every 6 hours over 144 hours under colistin concentrations ranging from 0 to 32 µg/mL. AUC metrics were calculated for growth (AUC_GR_), HR (AUC_HR_), and invasion (AUC_INV_). Color of boxes indicate either assay or experiment (gray), or bacterial strain or read out, where blue relates to the clinical isolate and green to invader.

### Media and Antibiotics

A colistin stock solution (10 mg/mL) was prepared by dissolving colistin sulphate powder (Cayman, USA) in deionized water followed by 0.2 µm filter sterilization and adjusting for purity and salt form. The solution was aliquoted and stored at -80°C for a maximum duration of three months and thawed on the bench on the day of the experiment. Sterile phosphate-buffered saline (PBS) was used for serial dilutions and daily preparation of 96-well plates containing antibiotics was automated using a benchtop liquid handler (OT-2, Opentrons). All experiments were carried out in Mueller-Hinton (MH) liquid and agar media (VWR chemicals). Mueller-Hinton powder (VWR, Belgium) was dissolved in water with or without agar and autoclaved. For agar plates containing colistin, the media was first cooled to 50°C then the colistin solution was added to reach the desired concentration before pouring the agar on the plates. Plates were stored at 4°C and used within a week.

### Strains and isolates

Ten colistin-susceptible clinical isolates of *P. aeruginosa* with varying levels of HR and minimal inhibitory concentration (MIC) variation (≤0.5 µg/mL) were selected from a large panel and used for this study(Lebreton et al. 2021). We generated a bioluminescent colistin resistant mutant to use as an invader when assessing invasion dynamics in co-cultures with the different clinical isolates. The ancestor strain was the colistin sensitive (MIC 0.5 µg/mL) *P. aeruginosa* PAO1-XEN41, harboring the constitutively expressed *luxCDABE* cassette for bioluminescence. To generate the resistant mutant, we serially transferred the strain in increasing concentrations of colistin until the strain could grow at 16 µg/mL (32x MIC).

### Growth and Antibiotic Susceptibility Determination

Colistin MIC and bacterial growth were determined using a microbroth dilution method following EUCAST guidelines(European Committee on Antimicrobial Susceptibility Testing 2003). Bacterial strains were streaked on agar plates and incubated overnight. After 24 hours, bacterial cells were harvested from the agar surface to prepare 0.5 McFarland standards in PBS (∼10^8^ colony-forming units [CFU]/mL), which were diluted to achieve a target inoculum of ∼10^5^ CFU/mL. After inoculation, 96-well plates were incubated with shaking at 150 rpm and grown at 37°C. Optical density (OD_600_) measurements to determine growth were taken hourly in a Fluostar microplate reader (Fluostar, BMG) over an 18-hour period. The final OD_600_ value was used to define MICs. All testing was conducted in triplicate over a colistin gradient between 0-8 µg/mL.

#### Quantification of growth

The Area Under the Curve (AUC) of the OD_600_ was used as a metric to quantify the growth (AUC_GR_) and calculated using the trapezoidal integration method shown in **Equation 1:**

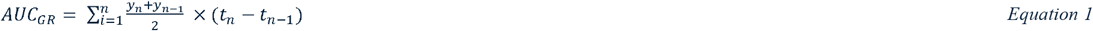

where *t* represents time (hours) and *y* corresponds to OD_600_ values at each time point, and the resulting AUC_GR_ values represent cumulative biomass at end of experiment, i.e. at 18 hours. This was performed for the drug-free condition as well as under colistin selection.

### Population analysis profiling

Population analysis profiling (PAP) was used to quantify the levels of colistin HR in the clinical isolates. Each isolate was cultured overnight in 25 mL MH broth. Serial dilutions (100 µL) of the overnight culture were plated on MH agar with colistin concentrations of 8, 16, and 32 µg/mL, and on antibiotic free plates. Sub-populations able to survive antibiotic concentrations above 8xMIC were considered heteroresistant(El-Halfawy and Valvano 2015). CFU were counted after 48 hours of incubation using an automated colony counting system (Acolyte, Synbiosis). The lower limit of detection was established at 1 colony for an undiluted sample. Data points below limit of detection was set to 1 CFU/mL.

#### Survival Frequency Determination

The survival frequency (F_survival_) was calculated per isolate and per antibiotic concentration used in the PAP. It was derived from the number of cells surviving at the colistin plate (N_R_) over the number of cells on the agar plate without antibiotic (N_T_), as shown in **Equation 2**.

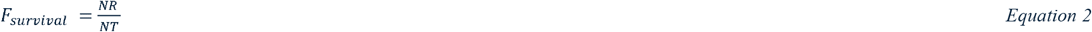

#### Quantification of Heteroresistance

To obtain a quantification of HR, the AUC of the PAP experiment was used as a metric (AUC_HR_). This was calculated in line with Equation 1, while instead of time *t* the colistin concentrations were used and instead of OD_600_ as *y* the log10(F_survival_) was used.

### Invasion dynamics into clinical isolates

We determined the invasion dynamics upon introducing the generated colistin-resistant *P. aeruginosa* strain, i.e. the invader, at low initial frequencies into cultures of the ten different clinical isolates (10^2^:10^6^). The low frequencies were used to simulate the invasion of a *de novo* resistance mutation or rare resistant migrant and represent the lowest quantifiable limit of detection. The invasion experiment was carried out for each clinical isolate across a colistin concentration gradient of 0 to 32 µg/mL and monitored over a period of 144 hours. At the start of the experiment each strain was grown overnight on MH agar and then used to prepare a 0.5 McFarland standard solution in PBS. These suspensions used to inoculate 10^2^ CFU/mL cell density of the invader with 10^6^ CFU/mL of each clinical isolate in 200 µL of MB broth in 96-well plates. Colistin concentrations were adjusted accordingly. Each condition was replicated six-fold. The cell density of the bioluminescent invader strain was monitored by measuring luminescence in relative light units (RLU) every 6 hours using a Fluostar microplate reader (Fluostar, BMG). Since RLU is highly correlated with OD_600_ and CFU/mL, it was used as a proxy for invader cell density(Koumans et al. 2024; Revvity 2023). To prevent nutrient depletion and extensive degradation of colistin, invasion cultures were serially passaged every 24 h, using a multi-blot replicator (VP 407, V & P Scientific), transferring 1.5 µL culture into freshly prepared MH media with the same colistin concentration.

#### Quantification of Invasion

The colistin-resistant invader strain was followed using luminescence, as outlined under the *Invasion dynamics into clinical isolates* section. To quantify the cumulative biomass of the invader, the AUC of the luminescence (AUC_INV_) was calculated aligned with Equation 1, where y instead represents RLU. The AUC_INV_ thus represents a summary metric of the invasion dynamics of the resistant strain over time. Final AUC_INV_ values are expressed in RLU·h, reflecting invader survival while in the presence of the clinical isolates and under the different colistin concentrations. For interpretation, we defined invasion success as reaching 6.4 RLU, i.e. the observed maximum.

### Software and data analysis

Data handling, data visualization, and statistical analyses were carried out in R 4.2.2.[R Core Team (2023)].

#### Statistical Analysis

To assess the variability in HR among the clinical isolates, a two-way ANOVA was performed. The ANOVA tested whether the mean log_10_(F_survival_) differed among isolates, with isolate identity and PAP-concentration as the independent variables and log_10_(F_survival_) as the dependent variable with a significance level of α = 0.01. Additionally, one way ANOVA’s were performed per PAP concentration, with isolate identity as the independent variables and log_10_(F_survival_) as the dependent variable (α = 0.01). To evaluate the strength and direction of the associations between the different derived AUC metrics (AUC_GR_, AUC_HR_, and AUC_INV_), Pearson correlation coefficients were calculated. Correlations were assessed for each AUC metric combination per colistin concentration using the mean AUC per isolate and condition. Only conditions with more than three mean data points were included.

## Results

The phenotypic characterization of the clinical isolates confirmed low MIC values (≤0.5 µg/mL, **Figure 2a**) and the respective full growth curves can be found in supplementary **Figure S1**. All experimental data can be found in the supplementary material.

**Figure 2.**
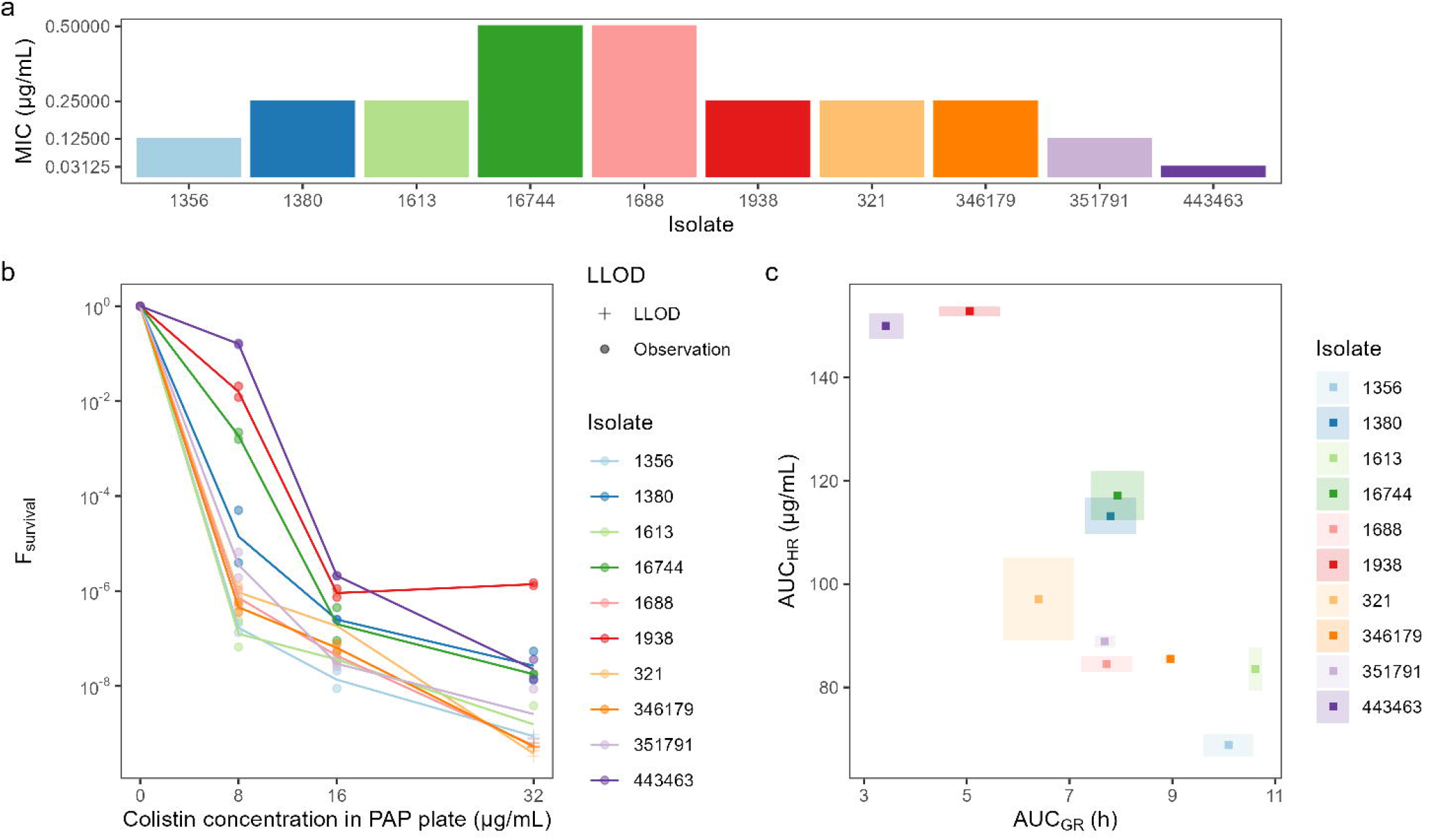
Characterization of clinical isolates. (a) MIC values (b) Heteroresistance profiles of clinical isolates measured by population analysis profiling (PAP) assay. Surviving fraction (F_survival_) across colistin concentrations (0–32 µg/mL). Points indicate an observation, and each line represents the mean of the replicates related to one isolate and concentration, colors indicate isolate identity, and plus marks indicate values below the lower limit of detection (LLOD: 1 CFU/mL). (c) Relationship between heteroresistance (AUC_HR_) and growth (AUC_GR_) in drug-free media. Each point represents the mean of one clinical isolate; shaded areas denote the range across replicates.

Despite low variation in MIC among clinical isolates, PAP analyses revealed differences in HR indicated by significant variability between isolates and within PAP plates on log_10_(F_survival_) (p < 0.01). Using a one-way ANOVA, we could confirm that the between-isolate variability was significant for all tested concentrations (p < 0.01). In the PAP analysis, F_survival_ declined with increasing colistin concentrations (**Figure 2b**). The mean log_10_(F_survival_) ± SD was -4.78 ± 2.16, -6.92 ± 0.725, -8.32 ± 1.12 for PAP concentration 8, 16, and 32 µg/mL, respectively. We also observed variation in growth across clinical isolates under antibiotic-free conditions, both in AUC_GR_ values (mean ± SD: 7.71 ± 2.19 h) (**Figure 2c**), and in carrying capacity, which were indicated by the final OD□□□ after 18 hours (mean ± SD: 1.07 ± 0.341) (**Figure S1, row 1**). We observed a strong negative correlation between AUC_GR_ in colistin-free conditions and AUC_HR_ (ρ = - 0.85, **Figure 2c and 3**). The correlation between AUC_GR_ and AUC_HR_ was less pronounced when investigating growth over the colistin gradient, and at 1-2xMIC it even showed a weak positive correlation (**Figure 3 and S2**).

**Figure 3.**
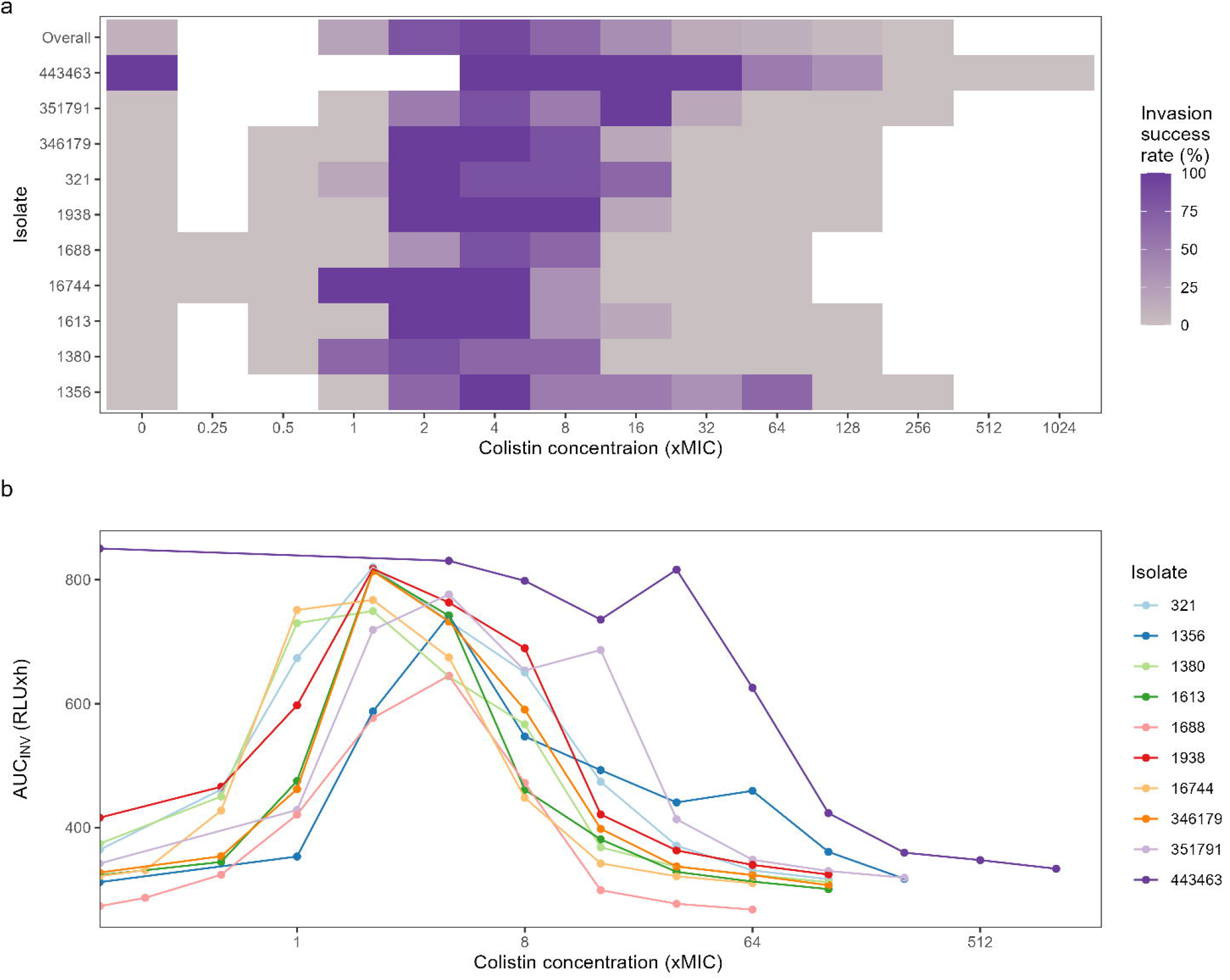
Pearson correlation coefficients (y-axis) comparing the mean growth (AUC_GR_), heteroresistance (AUC_HR_) and invasion (AUC_INV_) across colistin concentrations relative to MIC of the clinical isolate(x-axis). Error bars indicate the standard error, only correlations with more than 3 datapoints were included.

**Figure 3.**
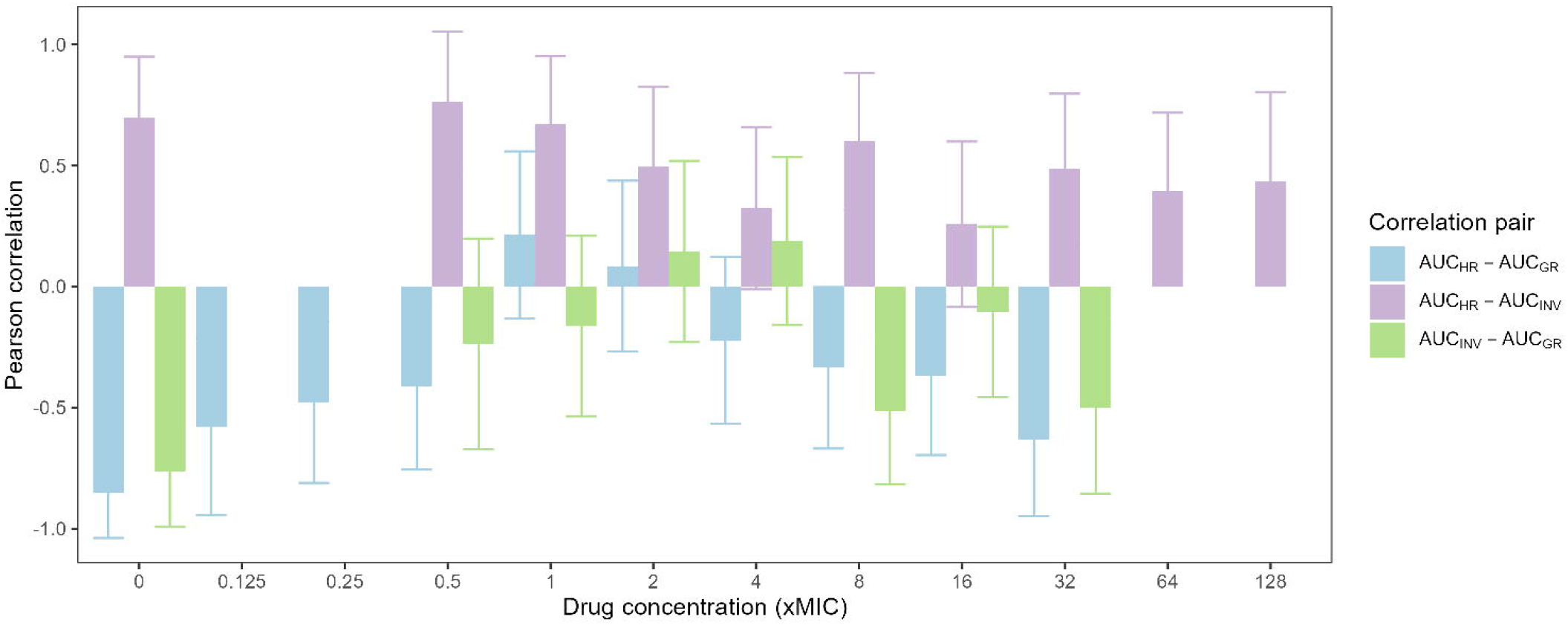
Panel (a) Invasion success rate per isolate (y-axis) and colistin concentration (x-axis). Colored tiles represent the invasion success while white tiles are non-tested conditions. The invasion success rate is calculated over six replicates. Panel (b) Relationship between colistin MIC (x-axis) and AUC_INV_ (y-axis) across isolates. Points represent mean AUC_INV_ values. Isolates are color-coded as shown in the legend.

### Invasion dynamics are shaped by growth, heteroresistance, and colistin

Invasion dynamics of the colistin-resistant (MIC 16 µg/mL) strain varied across clinical isolates and colistin concentrations (**Figure S3**). In the absence of colistin, the resistant invader failed to establish in 9 out of 10 co-cultures and the most consistent invasion success occurred at a colistin concentration of 4xMIC (**Figure 4a**). At sub-inhibitory concentrations of the clinical isolates (≤0.5×MIC), AUC_INV_ remained low and invasion success rate was low, suggesting that antibiotic selective pressure on the clinical isolates was insufficient to permit invader establishment (**Figure 4**). At intermediate concentrations (2-8×MIC), AUC_INV_ peaked in the presence of most isolates (**Figure 4b**). This range likely reflects a window where colistin suppressed the clinical isolates, thereby reducing competition of the co-cultured strains and allowing the resistant invader to survive and expand. The mean maximum AUC_INV_ across all co-cultures was 780 ± 58.7 RLU·h (**Figure S2**), representing a 2.16 time increase in AUC_INV_ compared to the colistin-free controls, on average. At higher concentrations (≥8xMIC), AUC_INV_ sharply declined across most co-cultures, consistent with widespread collapse of both susceptible clinical isolates and the resistant invader. This bell-shaped pattern, where AUC_INV_ peaked at intermediate antibiotic levels but diminished at both ends of the gradient, was observed across most clinical isolates (**Figure 4b**).

To better understand how growth dynamics of the clinical isolates shapes the invasion we investigated the correlation of AUC_GR_ and AUC_INV_. In a drug-free environment, we observed a negative correlation between AUC_GR_ and AUC_INV_ (ρ = - 0.76, **Figure 3, Figure S4**). When comparing the growth in colistin-free conditions and the invasion over the colistin gradient the correlation remained negative (ρ = -0.27 to -0.79, **Figure S5a)**. This correlation suggests that higher growth of the clinical isolate made it harder for the resistant strain to invade. This negative relationship persisted across most colistin concentrations (exception 4-8xMIC), though the magnitude of the correlation varied (range ρ: -0.16 to -0.51, **Figure 3**). Isolate 44343 exhibited different invasion dynamics than the rest of the isolates (**Figure 4**) and could potentially drive some of the investigated correlations. Therefore, we also examined correlations without isolate 44343 (**Figure S5**) and found less consistent results for concentrations >MIC.

Next, we determined whether HR is correlated with the invasion ability of the resistant strain by quantifying the relationship between AUC_HR_ and AUC_INV_ across colistin concentrations **(Figure 3 and S6)**. This analysis showed a consistent positive correlation between AUC_HR_ and AUC_INV_, which was strongest at sub-inhibitory concentrations (ρ > 0.65), and weaker but still positive at concentrations above the MIC (ρ = 0.26 - 0.60, **Figure 3**). Again, when determining the correlations in absence of isolate 44343, the results were less consistent and at higher concentrations (>8xMIC) there were instead negative correlations (**Figure S5b**).

## Discussion

This study examines the interactions between bacterial HR, growth, and antibiotic pressure in shaping the invasion dynamics of a laboratory-generated colistin-resistant *P. aeruginosa* strain. Using a set of clinical isolates with varying levels of HR but similarly low MICs, we created an invasion scenario in which the colistin-resistant strain must establish itself within a larger competitive population. By systematically analyzing phenotypic traits of the clinical isolates and their relationship with invasion dynamics, we observe that invasion is associated with both competitive exclusion and the presence of heteroresistant subpopulations, with selective pressure exerted by colistin further modulating these interactions.

Overall, we observed a negative correlation between AUC_GR_ and AUC_HR_, revealing a potential trade-off between growth capacity and phenotypic heterogeneity. In a colistin-free environment, clinical isolates with higher AUC_GR_ tended to display lower AUC_HR_, while isolates exhibiting greater AUC_HR_ tended to have lower AUC_GR_. This strong inverse relationship might reflect fitness costs associated with maintaining heteroresistant subpopulations linked to metabolic burden or regulatory instability(Andersson, Nicoloff, and Hjort 2019) or is a result of the higher fraction of slow-growing HR cells reduces the overall population growth. The correlation between AUC_GR_ and AUC_HR_ was less pronounced over the colistin gradient, and at 1-2xMIC it even showed a weak positive correlation, demonstrating how antibiotic pressure can modulate the association between these traits. From the perspective of the clinical isolates, these trade-offs could imply a tension between optimizing fitness to compete in drug-free conditions versus ensuring survival during antibiotic treatment.

In the context of invasion, isolates with low AUC_HR_ seem to be better at excluding resistant competitors as they display lower AUC_INV_ under no-drug conditions compared to isolates with high AUC_HR_. These findings suggest that a high AUC_HR_ could potentially facilitate invasion, although these results should be interpreted with caution, especially considering the high correlation of AUC_GR_ and AUC_HR_. Across clinical isolates, we observed this over the colistin gradient, particularly under low and moderate antibiotic pressure. However, the correlation seems to diminish with rising colistin concentration and was not retained during re-evaluation without isolate 443463 for most concentrations above 2xMIC. Consistent with the concept of the mutant selection window and mutant prevention concentration frameworks(Andersson and Hughes 2014), these results indicate that the selective advantage associated with heteroresistance is most pronounced under low and moderate antibiotic pressure, where both susceptible and less susceptible subpopulations can coexist. In these conditions, isolates displaying high AUC_HR_ supported greater invasion by the colistin-resistant strain. However, as the antibiotic concentration increased above MIC, this correlation weakened and was no longer evident in most conditions. This observation aligns with pharmacodynamic studies of colistin and polymyxin B showing that bacterial killing rates reach saturation above MIC concentrations, reducing fitness differences between competing subpopulations(Tam et al. 2005). Likewise, theoretical and empirical analyses of selective compartments(Baquero and Negri 1997) and heteroresistance dynamics(Levin et al. 2024) predict that once antibiotic concentrations exceed the range that permits partial survival of less susceptible cells, the ecological window favoring invasion closes, and selection among populations becomes neutral or uniformly lethal, as exhibited by our findings.

The correlation between AUC_INV_ and AUC_HR_ suggests that, beyond general population collapse, HR may influence the extent or timing of invader establishment. Overall, a positive correlation was observed over the colistin gradient. In this context, HR may reduce phenotypic uniformity within the clinical isolate population, by introducing survival heterogeneity among individual cells, and thus weaken competitive exclusion mechanisms. This could occur not only in drug-free environments but also under antibiotic selection. Notably, the observed correlations were sensitive to inclusion or exclusion of the clinical isolate with a deviant growth and HR profile (isolate 44343). When excluded, a negative correlation was observed at high concentrations, suggesting that the HR bacterial reservoirs that survive under high antibiotic selection can limit the invasion of the resistant strain. While our data does not directly capture bacterial sub-population during the invasion experiment, the observed correlation between AUC_HR_ and AUC_INV_ suggests that HR may contribute to shaping the ecological landscape in ways that affect the probability and magnitude of the invasion of the colistin resistant strain.

In antibiotic-free environments, invasion (AUC_INV_) was negatively correlated with the growth (AUC_GR_) of the clinical isolates, where AUC_INV_ reflected their ability to expand and compete for resources over time(Lenski et al. 2015). Clinical isolates with higher AUC_GR_ were less permissive to invasion by the colistin-resistant strain. This result is consistent with classical ecological theories of competitive exclusion(Lenski et al. 1998), and mirrors clonal interference patterns, where less-fit variants fail to expand due to resource monopolization by fitter lineages(Lenski 1991). In this study, the resistant invader acted analogously to such a variant and was outcompeted by the fitter population in the absence of antibiotic selection. This competitive landscape shifts with increasing colistin concentration as the resistant invader gains a fitness advantage and dominates the population. When high enough concentrations are achieved both subpopulations collapse, and no growth is observed. This is in line with the observed bell-shape invasion dynamics curve over the tested colistin concentrations, with an optimal invasion window around 2-4xMIC where colistin may have suppresses the dominant susceptible population while sparing the resistant invader.

Both clinical isolates with higher levels of phenotypic HR and isolates with lower levels of growth were associated with higher invasion. However, due to the strong negative correlation between HR and growth, especially in drug-free medium, it is not possible to disentangle what is the main driving force shaping the invasion dynamics. Our results indicate that HR and growth may act as opposing ecological forces under antibiotic pressure. While growth confers a competitive advantage in drug-free environments, it may reduce plasticity under drug treatment. This is also indicated by the persistent negative correlations between drug-free growth and invasion over the colistin gradient. In contrast, heteroresistant populations, despite growth costs, can persist and create conditions that favor the establishment of resistant variants when colistin is present. The observed isolate-specific variation in HR highlights the potential for differential evolutionary trajectories, were fitter isolates limit invasion through competitive exclusion, while heteroresistant populations, despite their growth costs, may serve as reservoirs for resistance evolution under antibiotic pressures.

## Conclusion

This study has shown that growth, HR, and antibiotic pressure seem to influence the invasion dynamics of colistin-resistant *P. aeruginosa*. Understanding these factors interactions is critical for predicting resistance outcomes and for designing treatment strategies that minimize ecological opportunities for resistant strains to establish and expand. Importantly, such understanding cannot be obtained solely on MIC susceptibility testing but requires a deeper phenotypic characterization as demonstrated here.

