## Supplemental material for "The influence of heteroresistance, growth and antibiotic selection in shaping the invasion dynamics of colistin resistant *Pseudomonas aeruginosa*"

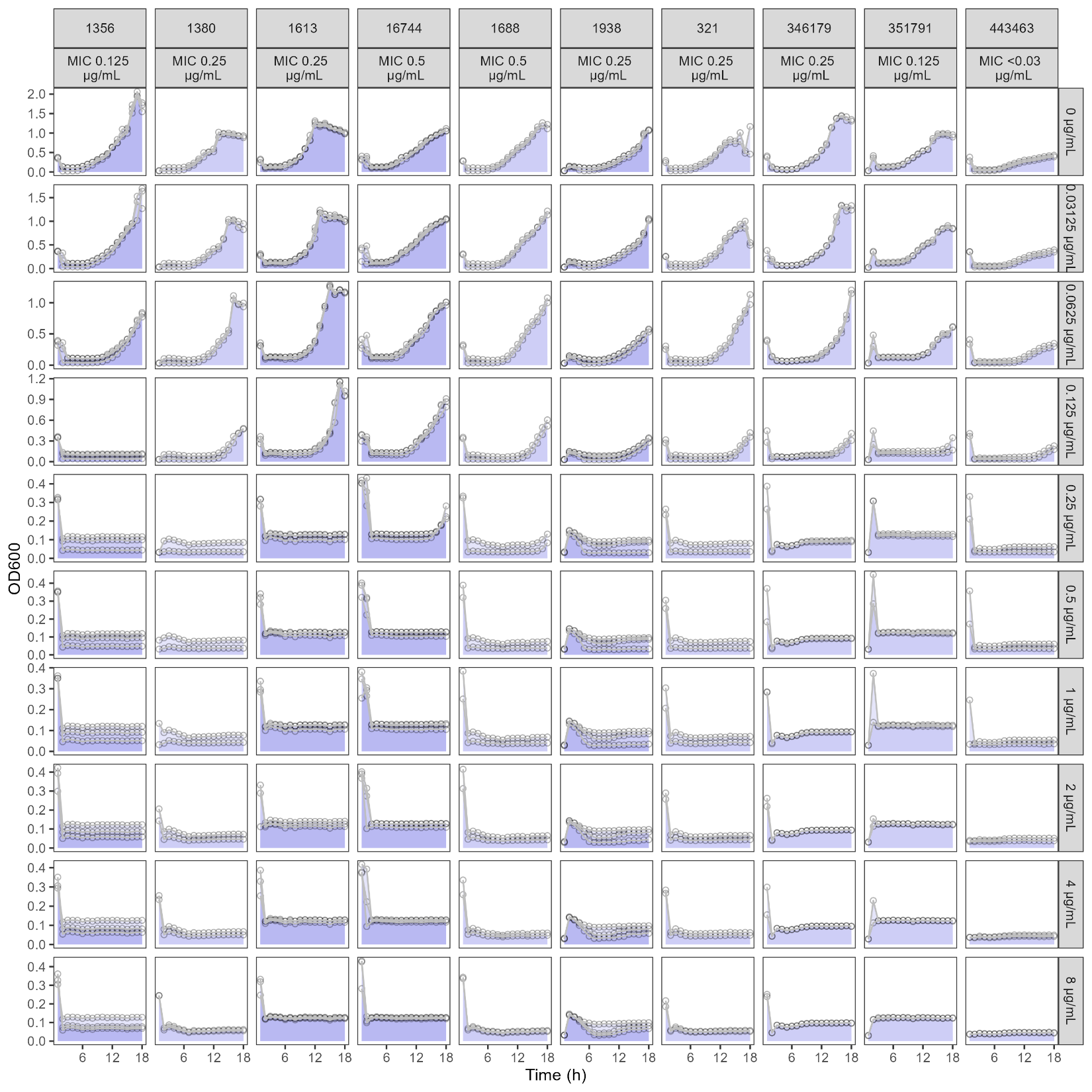
***Figure S1.*** Growth dynamics of the clinical isolates (columns) over a colistin gradient (rows), measured as OD600 over 18 hours. The shaded area represents the AUC (Area Under the Curve), quantifying growth under each condition.

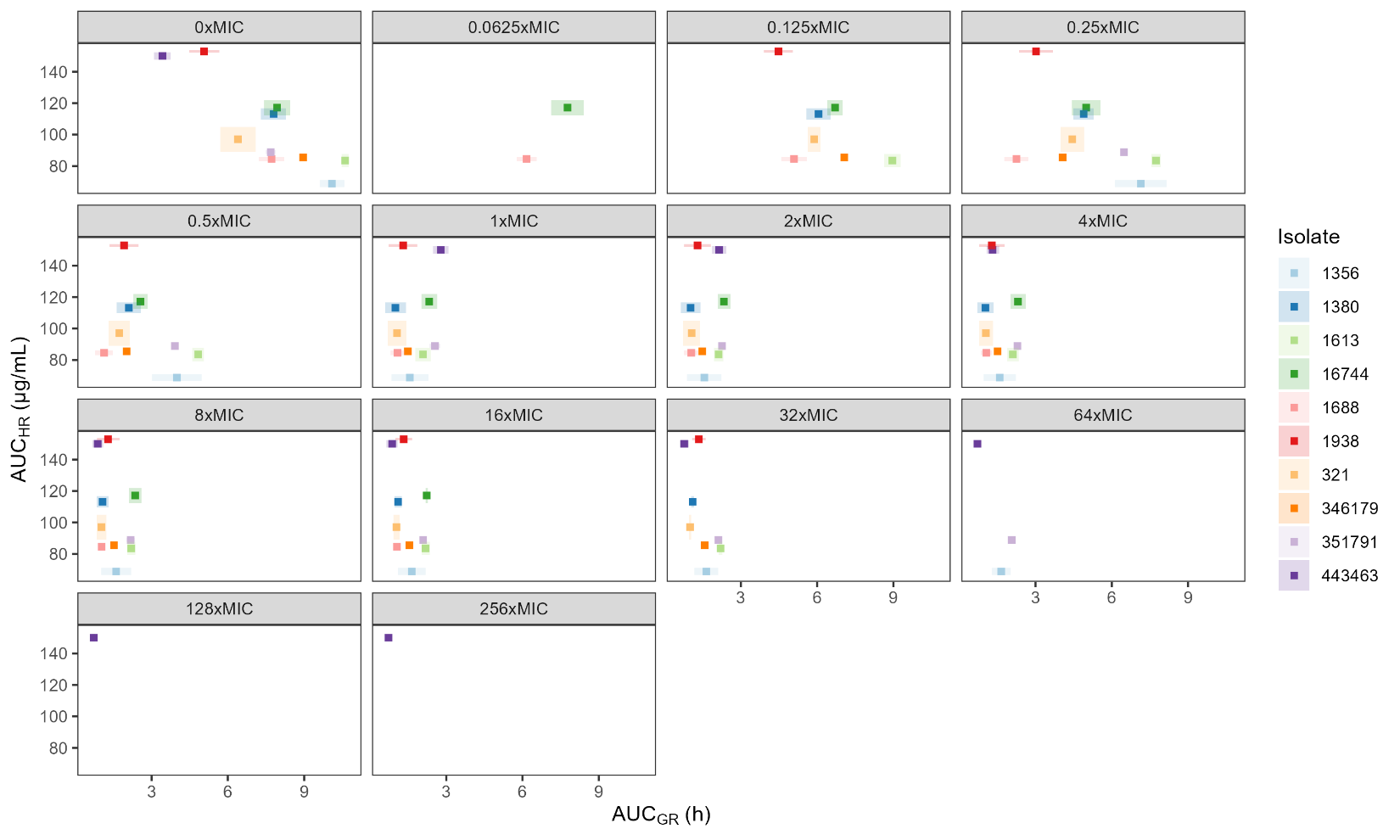

**Figure S2.** Relationship between growth (AUC_GR_) and heteroresistance (AUC_HR_) across colistin concentrations. Each point represents the mean of one clinical isolate; shaded area denotes the range across replicates.

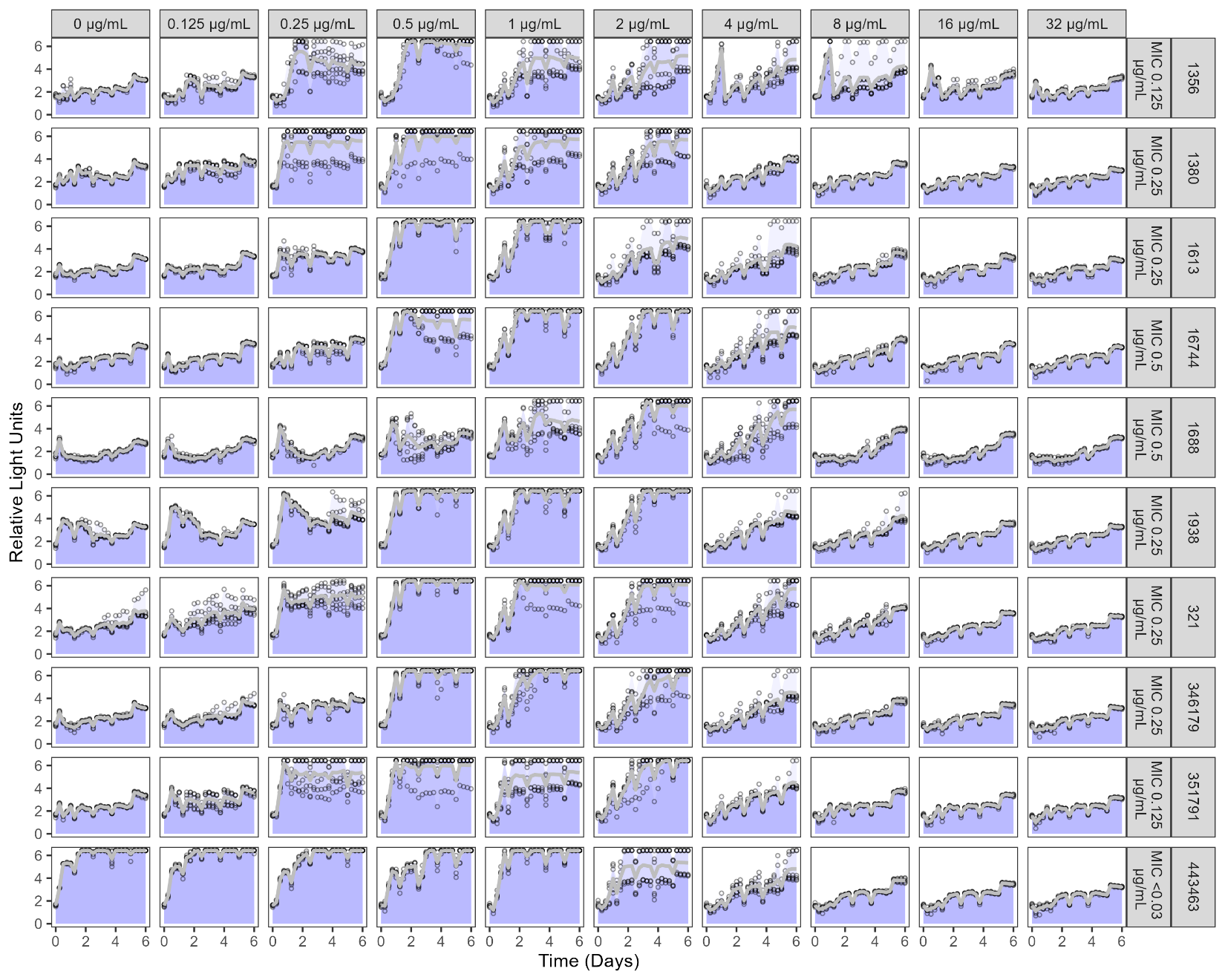

***Figure S3****.* Invasion profiles of 10 clinical isolates across colistin concentrations. Invasion dynamics of the colistin-resistant invader in co-culture with 10 clinical isolates, each labeled by their corresponding MIC (row labels), across increasing colistin concentrations (column labels). Luminescence (Relative Light Units) was measured over 6 days to track invader survival. The shaded area represents the AUC (Area Under the Curve), quantifying invasion under each condition.

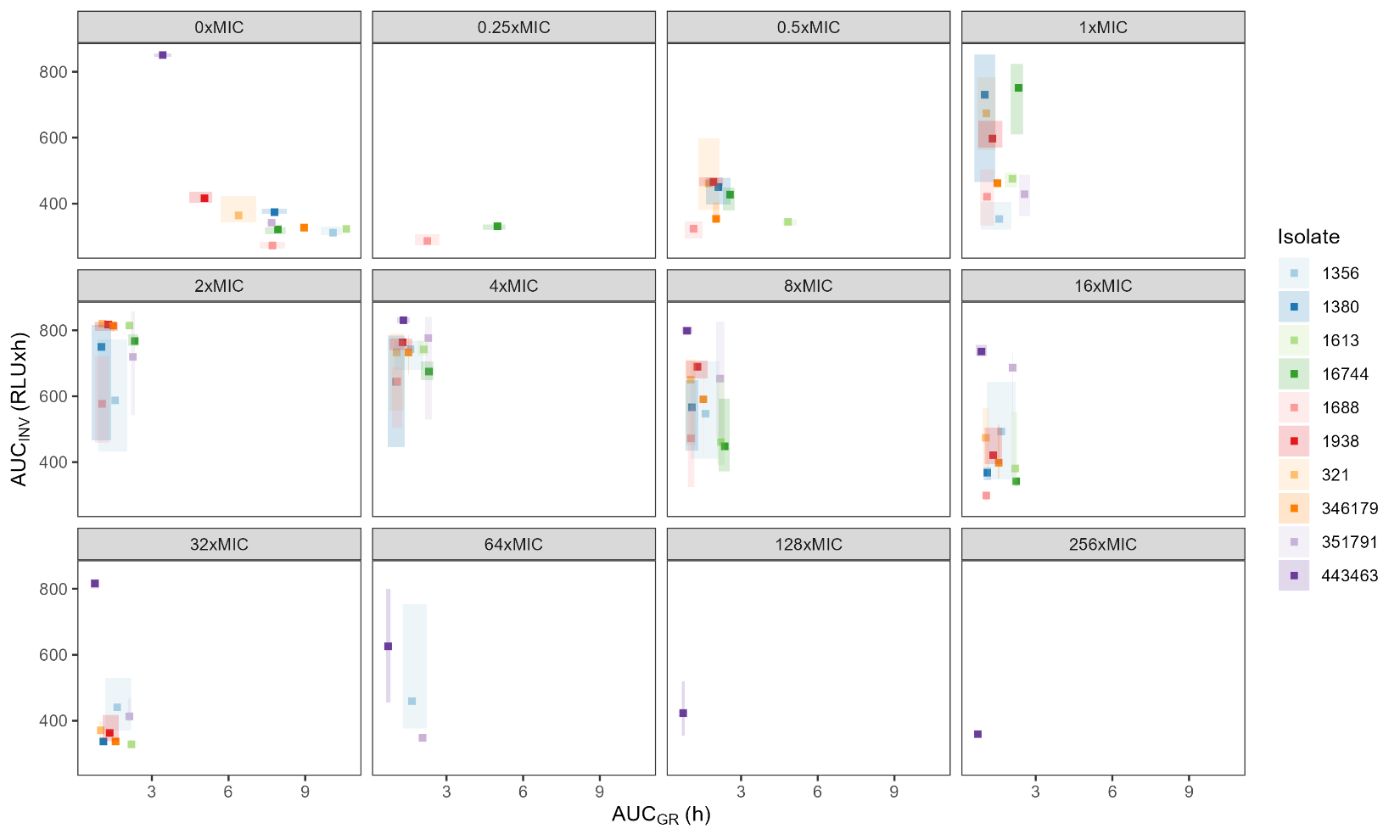

**Figure S4.** Relationship between growth (AUC_GR_) and invasion (AUC_INV_) across colistin concentrations. Each point represents the mean of one clinical isolate; shaded area denotes the range across replicates.

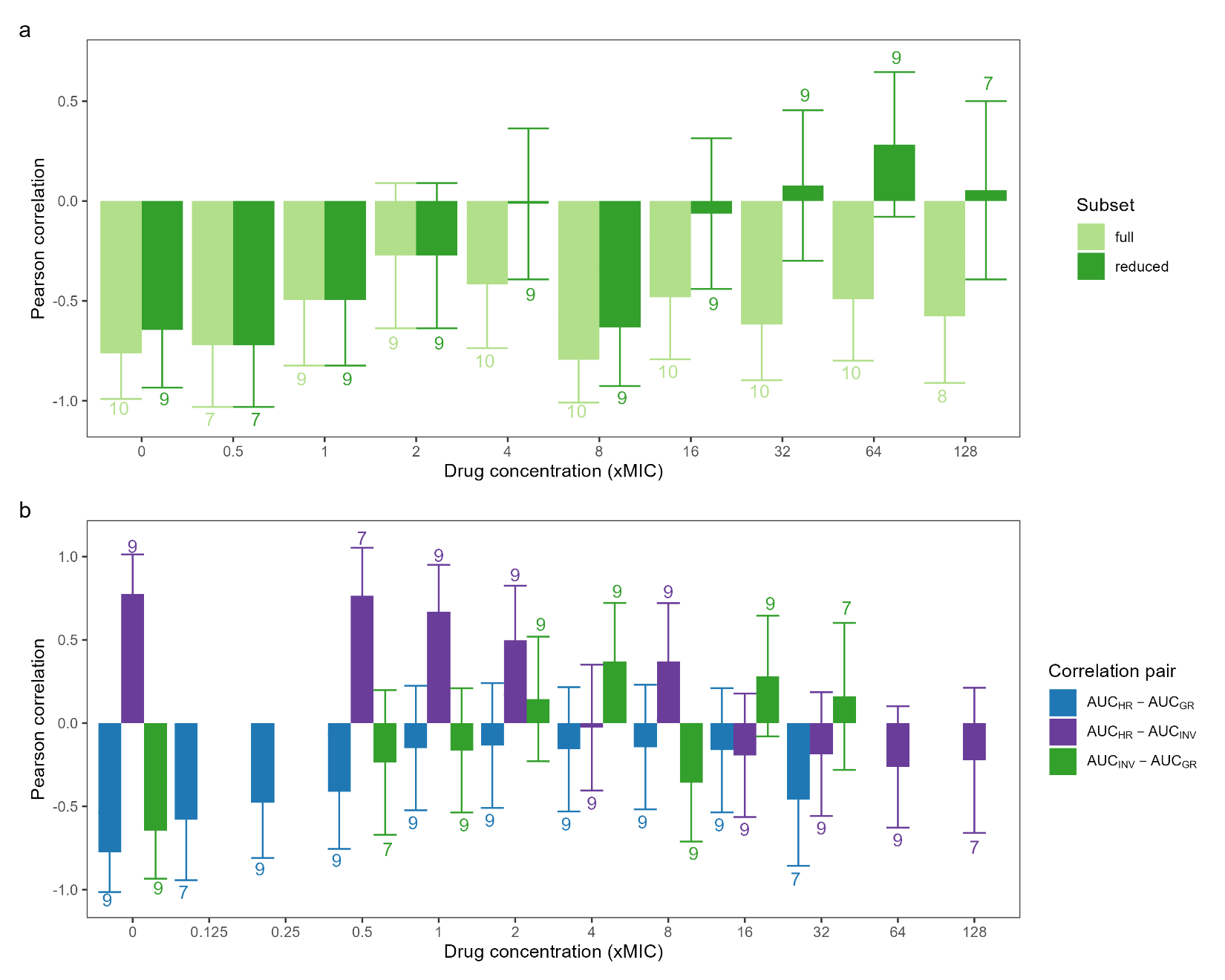

**Figure S5.** Pearson correlation coefficients (y-axis) comparing the mean AUC metrics over a colistin gradient relative to MIC (x-axis). Values show number of datapoint underlying the correlation and error bars indicate the standard error, only correlations with more than 3 datapoints were included. a) correlation between growth (AUC_GR_) and invasion (AUC_INV_) for the full data set or the reduced where isolate 443463 is excluded. b) correlations between AUC_GR_, heteroresistance (AUC_HR_) and AUC_INV_ for the reduced data set.

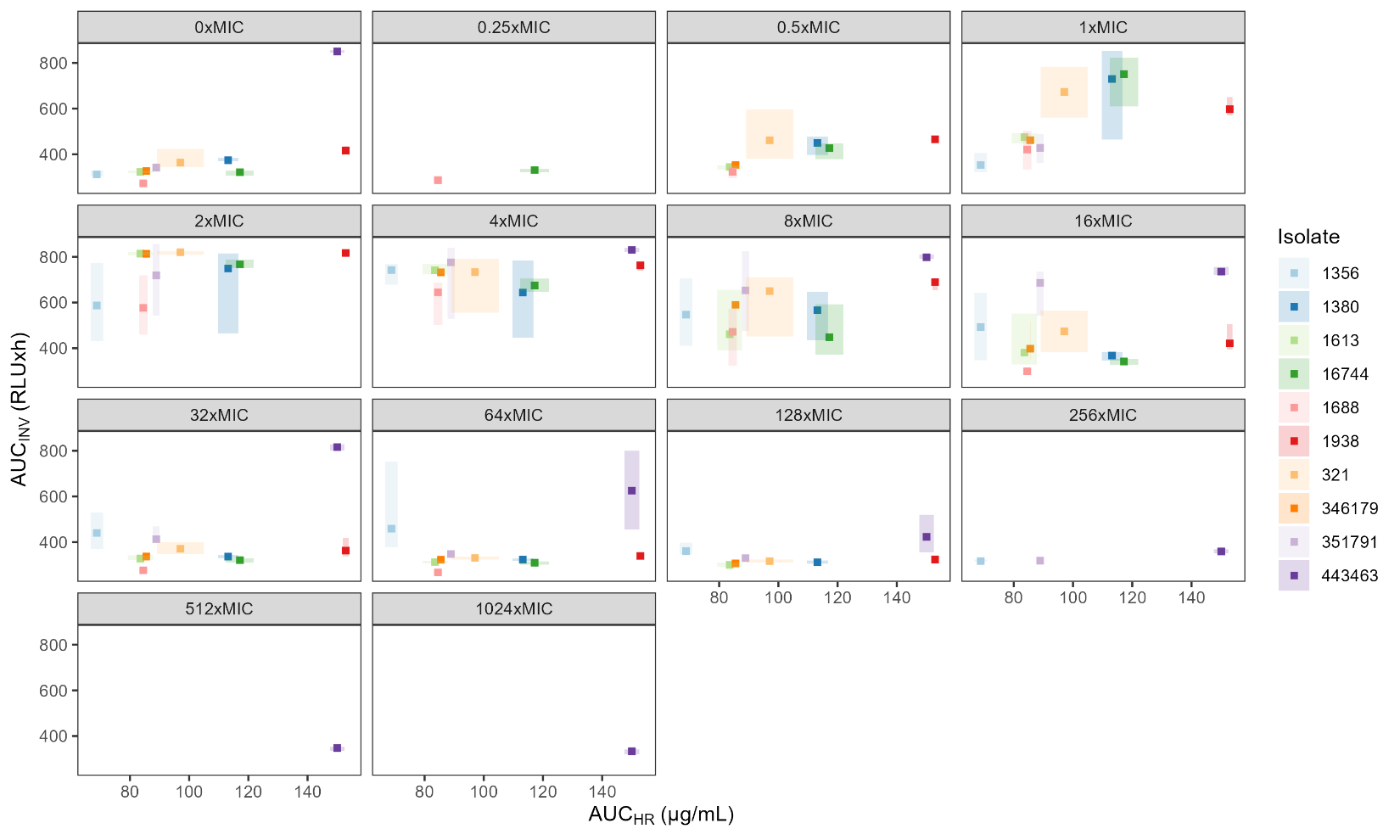

**Figure S6.** Relationship between heteroresistance (AUC_HR_) and invasion (AUC_INV_) across colistin concentrations. Each point represents the mean of one clinical isolate; shaded area denotes the range across replicates.
